# Data-driven filtration and segmentation of mesoscale neural dynamics

**DOI:** 10.1101/2020.12.30.424865

**Authors:** Sydney C. Weiser, Brian R. Mullen, Desiderio Ascencio, James B. Ackman

## Abstract

Recording neuronal group activity across the cortical hemispheres from awake, behaving mice is essential for understanding information flow across cerebral networks. Video recordings of cerebral function comes with challenges, including optical and movement-associated vessel artifacts, and limited references for time series extraction. Here we present a data-driven workflow that isolates artifacts from calcium activity patterns, and segments independent functional units across the cortical surface. Independent Component Analysis utilizes the statistical interdependence of pixel activation to completely unmix signals from background noise, given sufficient spatial and temporal samples. We also utilize isolated signal components to produce segmentations of the cortical surface, unique to each individual’s functional patterning. Time series extraction from these maps maximally represent the underlying signal in a highly compressed format. These improved techniques for data pre-processing, spatial segmentation, and time series extraction result in optimal signals for further analysis.

## Introduction

Optical techniques have long been used to monitor the functional dynamics in sets of neuronal elements ranging from isolated crustacean nerve fibers^1^ to whole regions of mammalian cerebral cortex^2, 3^. Imaging calcium flux with calcium sensors ^4, 5^ allows monitoring of neural activity within populations across the entire neocortex with high enough spatiotemporal resolution to monitor sub-areal networks of the neocortex ^6, 7^. These techniques have the potential to map group function at unprecendented resolution and scale across the neocortical sheet, however computational methods for analyzing these datasets are underdeveloped. An understanding of cerebral dynamics at multiple scales is important for exploring how environmental and genetic influences give rise to altered neural connectivity patterns linked to behavioral phenotypes.^8, 9^

Wide-field imaging of neuronal calcium flux can offer mesoscale observation of cortical neural dynamics, providing a view of supracellular group organization intermediate to that of microscale (cell) and macroscale (tissue lobe) investigations– however it is affected by issues common to topical imaging recordings. Body or facial movements can create large fluctuations in autofluorescence of the brain and blood vessels,^10^ which produce significant artifacts in the data. These techniques offer corrections of signal from the absorbance of GCaMP fluorescence by hemoglobin, but do not address the combination of movement, vessel artifacts, and surface optical aberrations.

Eigendecompositions can be used to identify and filter components of signal,^11–13^ and present a flexible method of filtering that is not hardware dependent, and can be applied to any video dataset regardless of the recording hardware or parameters. Independent Component Analysis (ICA)\cite{[@article]{hyvarinen independent 2000} has been previously applied to fMRI and EEG data with varying success; identifying intrinsic connectivity networks, rather than identification of individual areas, and artifacts that represent large-scale effects, rather than spatially localized effects.^14–18^ We hypothesize that this is due to the lower density of spatial sampling in fMRI and EEG data. Wide-field calcium imaging provides a unique combination of spatially and temporally resolved dynamics across the cortical surface, with scale ranging from complex activation patterns in high-order circuits, to discrete activations hundreds of microns in diameter, to whole cortical lobe activity patterns.^6, 7^ Researchers have recorded wide field calcium dynamics at frame rates ranging from 5-100Hz.^6, 19, 20^ In addition, spatial resolution varies between different researchers’ setups, but is typically in the range of 256×256 to 512×512 pixels (0.06 to 0.2 megapixels) for the entire cortical surface, and is often further spatially reduced for processing.^6, 19, 21^ Selection of resolution is often dependent on the video observer’s perceived quality of the data or available computational resources, rather than a quantified comparison of signal content.

It is common to use sensory stimulation to identify specific regions in the neocortex, and align a reference map based on the location of these defined regions.^21–23^ Even if these maps are reliable for the location of primary sensory areas, they often lack specificity for higher order areas, or even completely lack sub-regional divisions. This is especially true in areas with a high degree of interconnectedness, with overlapping functionality, such as motor cortex.^24^ Moreover, there is evidence that the shape and location of higher order regions can vary from subject to subject.^25, 26^ Improper map alignment or misinformed regional boundaries can lead to a loss in dynamic range between signals across a regional border. Thus, to extract the most information from a recorded dataset, the level of parcellation must reflect the quality and sources present within the data. Thus a flexible data-driven method is necessary that respects functional boundaries of the cortex and is sensitive to age, genotype and individual variation.

Here we present an ICA-based workflow that isolates and filters artifacts from calcium imaging videos. We explore the resolution-dependent effect on ICA quality of signal extraction, and find a quantified increase in ICA signal separation for collecting wide-field calcium imaging at mesoscale resolution. Using neural components, we additionally generate data-driven maps that are specific to functional borders from individual animals. We further use these maps to extract time series from functional regions of the cortex, and show that this method for time series extraction produces a reduced set of time series while optimally representing the underlying signal and variation from the original dataset. Together, these methods provide a set of optimized techniques for enhanced filtering, segmentation, and time series extraction for wide-field calcium imaging videos.

## Results

To record neural activity patterns, we transcranially image fluorescence from a genetically encoded calcium indicator across the entire mouse neocortex. The genetically encoded calcium indicator, GCaMP6s, is expressed in all neurons under the control of the Snap25 promoter. We expose and illuminate the cranium with blue wavelength light and capture fluoresced green light with a sCMOS camera at high spatial resolution (2160×2560 pixels, 5.5 megapixels; ∼6.9 µm/pixel). To observe the spatiotemporal properties of these population neural activity patterns, we crop the video to only neural tissue, and compare the change in fluorescence over the mean fluorescence: ΔF/F (fig. 1a,b).

**Figure 1:**
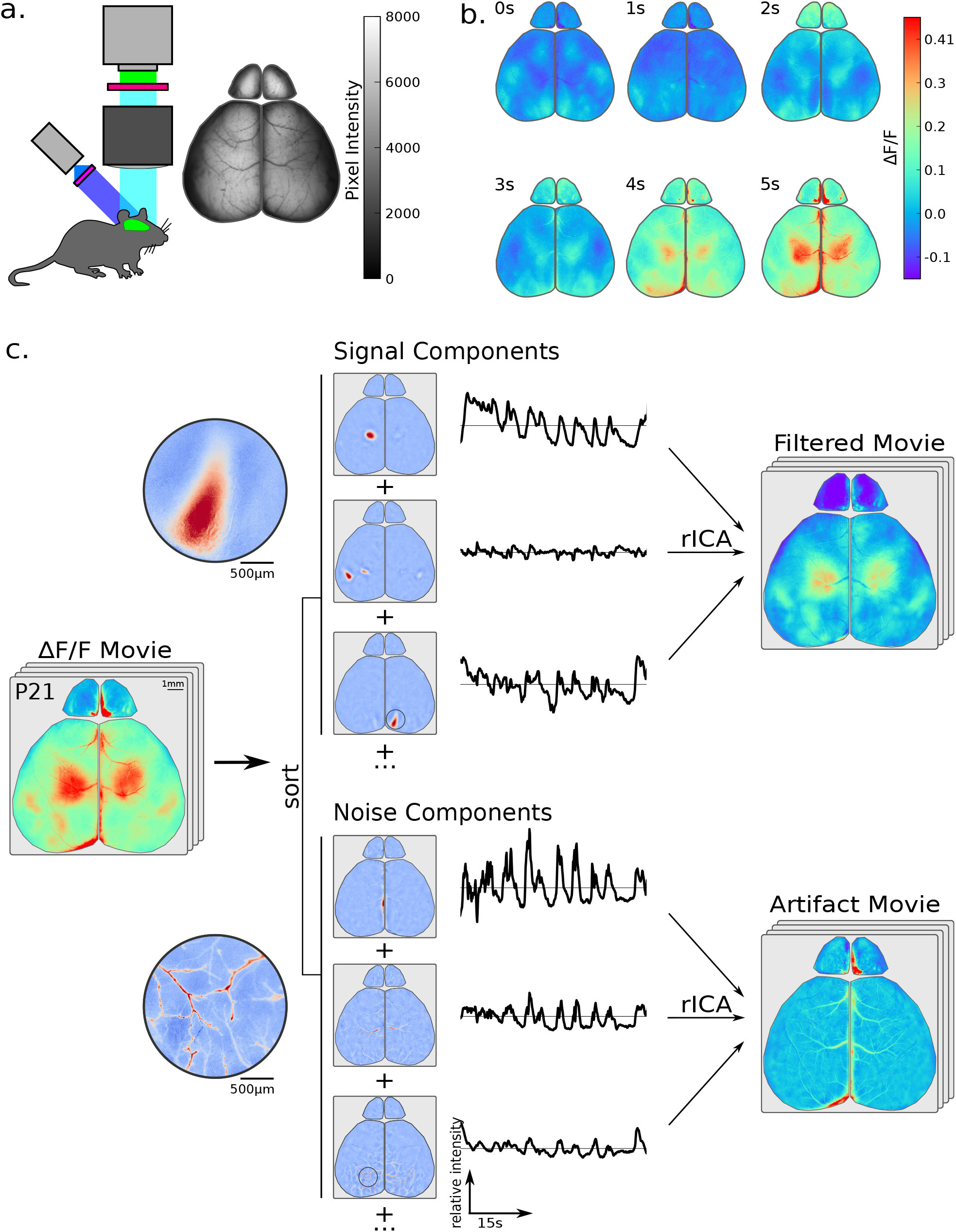
Transcranial calcium imaging video data is separated into its underlying signal and artifact components, and can be rebuilt from only signal components for artifact filtration. a) Recording schematic and fluorescence image of transcranial calcium imaging preparation, cropped to cortical regions of interest. b) Sample video montage of raw video frames after dF/F calculation. c) Independent Component Analysis (ICA) video decomposition schematic. A dF/F movie is decomposed into a series of statistically independent components that are either neural, artifact, or noise associated (not displayed). Each component has an associated time course from the ICA mixing matrix. Signal components can be rebuilt into a filtered movie (rICA). Alternatively, artifact components can be rebuilt into an artifact movie. Circular panels show higher resolution spatial structure in bottom example components.

### ICA Filtration

A spatial ICA decomposition on a de-meaned video produces a series of spatial components and a mixing matrix, representative of the component’s influence at each frame in the video (fig. 1c). The components are sorted by influence over the temporal variance, and flipped so that they all represent positive effects. The independent components can be sorted into 3 major categories based on their spatiotemporal properties: neural components, artifact components, and noise components (not shown).

Signal components represent a distinct area of cortical tissue, which we refer to as it’s cortical domain. The spatial morphology of these signal components can vary in both spatial extent and eccentricity. Occasionally signal components can also contain a multiple domains, with similar enough activation patterns to be identified as a single neural component. In the example, the second signal component appears to represent a higher order visual network, with multiple domains on the left hemisphere, and a small mirrored domain on the right hemisphere.

Artifact components can take many forms, including various blood vessels, movement artifacts, optical surface artifacts, etc.^27^ The top two artifact examples likely represent hemodynamics from the superior saggital sinus vein with the bottom artifact likely represents blood flow through the middle cerebral artery.^28^ A very high resolution map of the vessel patterns can potentially be rebuilt from these components, with branching structures as small as 12 µm in diameter (shown above in zoom panel). Noise components lack a spatial domain, and have little to no temporal structure. Signal and artifact components can be sorted manually in graphical user interface (fig. S1) or with a machine learning classifier.^27^

Video data can be reconstructed using any combination of these components. In particular, a filtered video can be constructed by excluding all artifact components. Since there was a low-frequency hemodynamic effect common to GFP control mice, we re-added the mean signal filtered by a 0.5 Hz wavelet high pass filter.^27^ The noise components can be additionally excluded for denoising video data with high spatial noise. For our analyses, we did not exclude ICA noise components when rebuilding. The artifact movie can also be reconstructed to verify that desired signal was not removed with the artifact filtration (fig. S4 (Video)).

### Noise Sorting and Resolution Analysis

Non-noise components can be separated from noise components from their visual differences, as well as from their log temporal variances, or lag-1 autocorrelations. Non-noise components have spatial structure and a high lag-1 autocorrelation, corresponding to waves of structured calcium activity in a specific location. Conversely, noise components are highly dispersed across the cortex, and have a low lag-1 autocorrelation. The lag-1 autocorrelation metrics from these two groups are so polarized, that it is straightforward to separate these populations by their lag-1 autocorrelation alone (fig. 2a).

**Figure 2:**
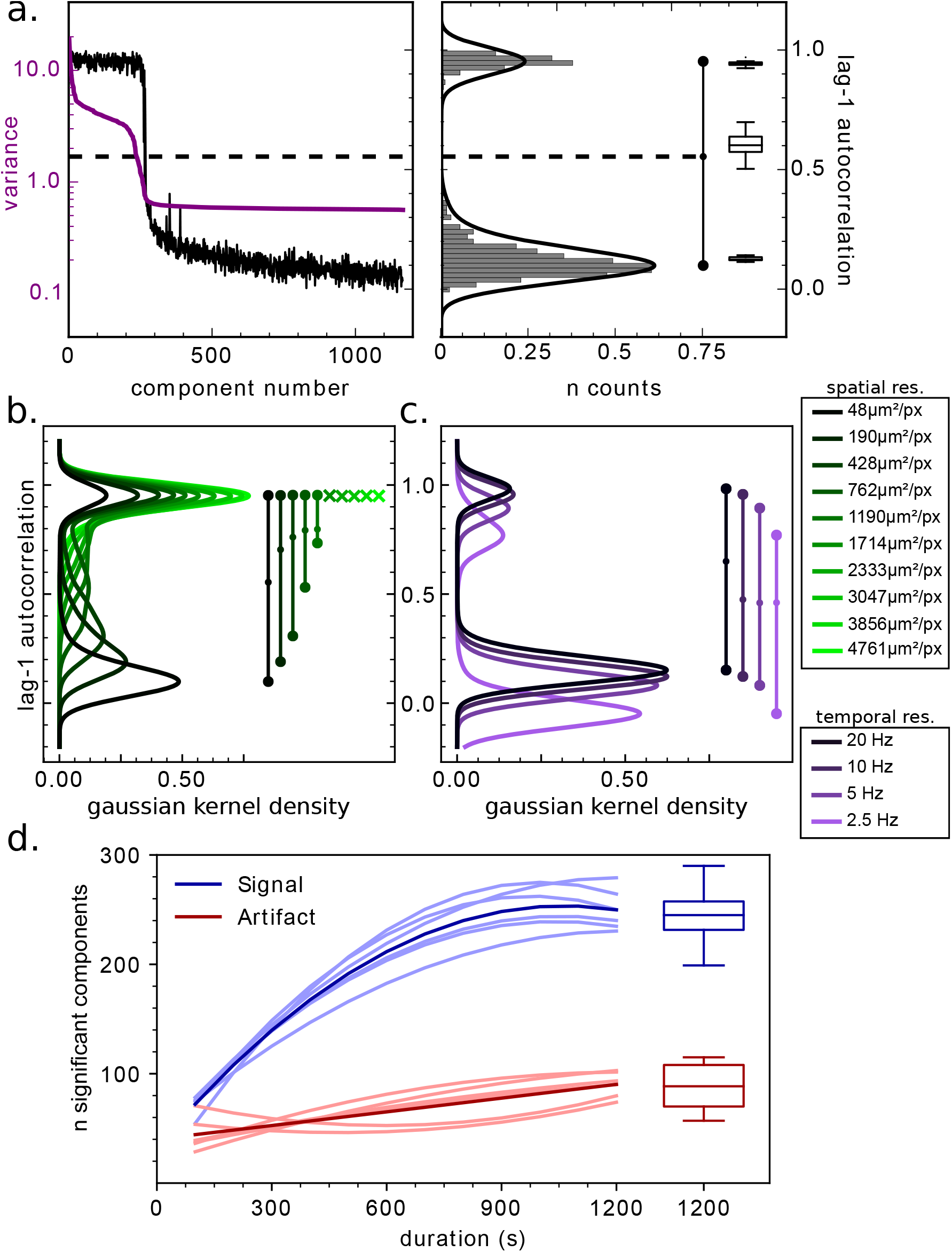
ICA decomposition quality is sensitive to recording duration, spatial and temporal resolution. a) Distributions for lag-1 autocorrelation (black) and temporal variance (purple) are displayed for components 1-1200. A dotted line representing the cutoff determined from the distribution in the right panel. In the right panel, a horizontal histogram on the lag-1 autocorrelation with a two-peaked kernel density estimator (KDE) fit reveals a two peaked-histogram, summarized by a barbell line. Group data for each peak, as well as the central cutoff value is summarized by the boxplots on the right (n=16 videos; from 8 different animals). b) 2-peaked KDE fits of horizontal histogram distributions under various spatial downsampling conditions, with barbell summary lines on the right. After spatial resolution decreases beyond 1714 µm^2^/px, this two peak structure collapses, and an x denotes the primary histogram peak. c) 2-peaked KDE fits of horizontal histogram distributions under various temporal downsampling conditions, with barbell summary lines on the right. d) Component stabilization for different length video subsets of six 20-minute video samples. (n=6 videos from 3 different animals) Individual thin lines show polynomial fit to signal or artifact components under each time condition. Thick lines denote the curve fit of the mean number of components in each category across these six experiments. The group distribution of components at 20 minutes is summarized by the boxplot on the right (n=16 videos; from 8 different animals).

To automate this sorting process, a two-peak kernel density estimator (KDE) was fit to the histogram of lag-1 autocorrelation data. The KDE distribution is an easy way to summarize the two major peaks, as well as the minima between them, defined as the noise cutoff. The locations of these peaks, and the minima between them is highly stable across our 8 P21 test recordings. We found the non-noise peak (*p*1) at an autocorrelation of 0.94 ± 0.01, and a noise peak (*p*2) at 0.13 ± 0.01. The central cutoff minima was slightly more variable, with an autocorrelation value of 0.61 ± 0.05. A high degree of separation between these peaks (*d*_*p*−*p*_ = 0.82 ± 0.01; *p* < 0.001) suggests that the signal and noise signal sources were completely separated, and thus all signal sources were distinctly identified.

To test how ICA component separation is affected by spatiotemporal resolution and video duration, we altered properties of the input video and observed its effects on the quality of signal separation through lag-1 autocorrelation distributions. Reducing the spatial resolution resulted in a steady decrease in peak separation, until the dual peaked structure collapsed at a resolution of 1714 µm/px (fig. 2b).

Increasing the sampling rate above 10Hz showed little to no effect on the peak to peak distance (Δ_*p*−*p*_ < 0.01), and a slight decrease in the autocorrelation of the primary peak (Δ_*p*1_ = 0.03), but temporal downsampling below 10Hz resulted in a shifting of the signal and noise peaks (Δ_*p*1_ = 0.06), and a reduction in the peak to peak distance (Δ_*p*−*p*_ = 0.02). This result agrees with previous analyses finding 10Hz as the maximal sampling frequency required for measuring population calcium dynamics.^22^ Together, these findings suggests that the separation quality of our captured dynamics are highly sensitive to spatial resolution, and not as sensitive to temporal resolution. We considered collecting spatial samples higher than our current resolution of ∼6.9 µm/px, but computing decompositions on datasets this large would push the limits of available computing.

To determine the ideal duration of video collected, we calculated the number of significant signal and noise components for various video durations. We found that for ICA decompositions on an activity patterns from a P21 mouse, the number of neural and artifact components leveled off after 20 minutes. Population analyses showed that this number was highly similar among P21 mice (n signal components: 244 ± 25.7; n artifact components: 87.2 ± 20.7).

### Generating Domain Maps

In addition to their applications for filtering, the components also are a rich source of information about spatial distributions of signal within the cortex. Components across the cortex show a wide diversity of spatial characteristics, and represent a detected independent unit of signal. We use the spatial domain footprints of each signal component to create a data-driven ‘domain map’ of the cortical surface by taking a maximum projection through each component layer (fig. 3a). For analysis, 8 maps were created, with an average of 230 ± 14 detected domains. Domains were then manually sorted into regions, with informed decisions based on network analysis, metric comparisons, and reference maps (fig. S3).

**Figure 3:**
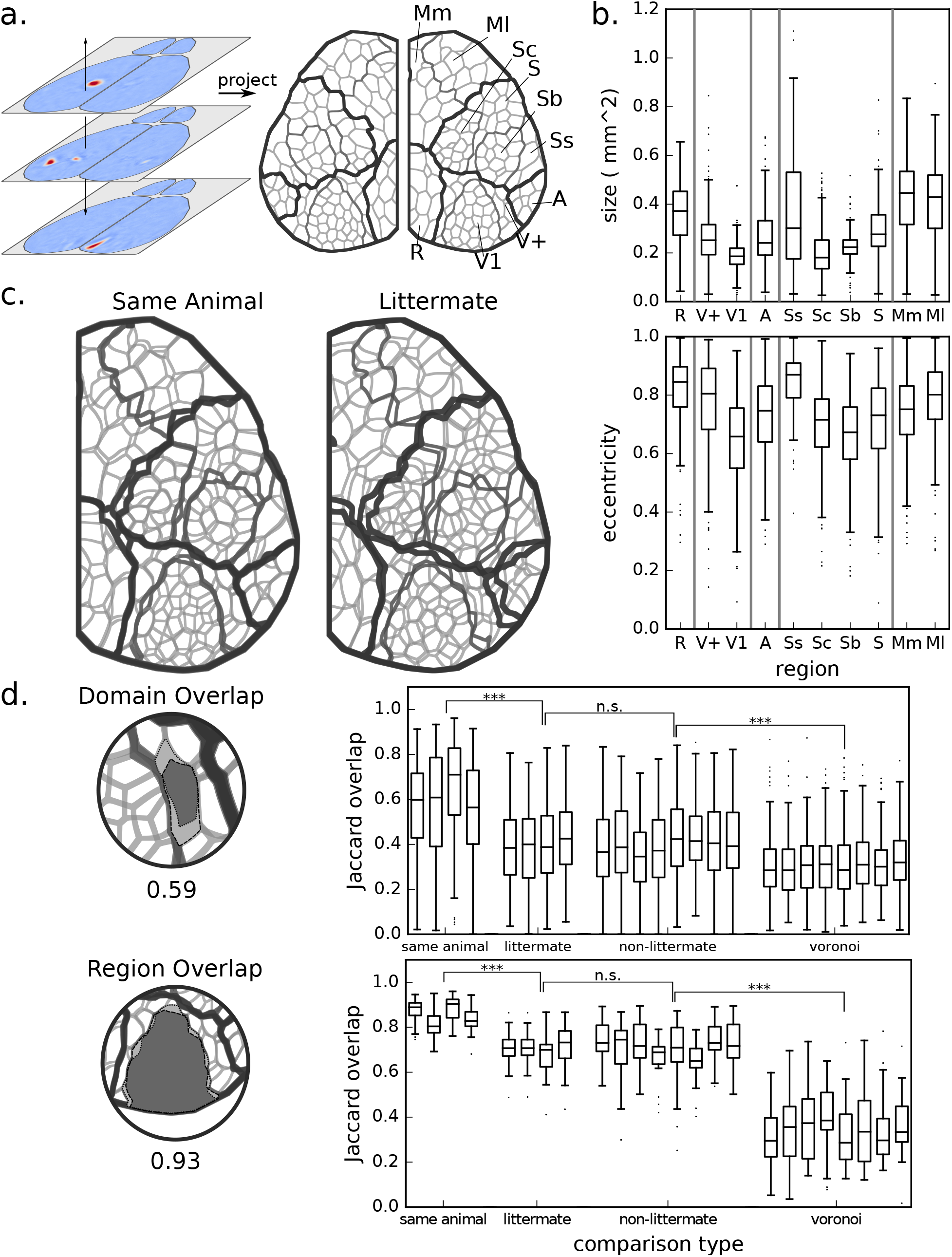
Domain maps are created from ICA components and are unique to each recording, but highly similar among individual animals. a) Schematic of domain map creation. A maximum projection is taken through each blurred signal component to form a domain map. Cortical domains are assigned to one of 10 identified regions, with region divisions denoted by the bold lines. b) Domain area and eccentricity by region. Population analysis of distribution of spatial characteristics individual domains within defined regions across multiple recordings (n=16 videos; from 8 different animals). c) Example overlay of one domain map on another from the same animal. Individual domain or region overlap is calculated using the Jaccard index (intersect / union). d) Population analysis of the Jaccard index for domain and region overlap comparisons. Maps are generated from a different recording on the same animal, a littermate, a non-littermate, or a randomly generated voronoi map. Significance is calculated using a two-way ANOVA, followed by post-hoc t-test analysis with Holm-Sidak corrections.

Domains did not exhibit uniform spatial characteristics across the neocortex. Different detected regions have different spatial characteristics such as area (ANOVA F=139, *p* < 0.001), as well as eccentricity (ANOVA F=60.0, *p* < 0.001). Generally, higher order and motor regions (R, V+, Ss, Mm, Ml) had larger domains than primary sensory areas (V1, A, Sc, Sb, S) (p>|t| = 0.000), and also exhibited higher eccentricity (p>|t| = 0.000).

To test the meaning of these maps, a series of comparisons were performed. Pairs of maps were 5 overlayed on top of each other (fig. 3c-d), and every domain was compared to its nearest domain in the comparison map. The Jaccard overlap was calculated for each of these domain pairs, and quantified for each pair of map comparisons. For a null hypothesis, randomly generated Voronoi maps were also compared.

Maps generated from different recordings from the same animal were found to be highly overlapping, and hence more similar (fig. 3d, top; *p* < 0.001). There was no significant difference in comparisons between littermates vs non littermates. Non-littermate map comparisons were significantly more similar to each other than to voronoi maps (*p* < 0.001).

We additionally quantified whether detected regions were similar across map comparisons. We again found that comparison between maps from the same animal were highly similar (fig. 3d, bottom; *p* < 0.001), no difference was found between littermates and non-littermates, and comparisons between different animals were significantly more similar than a comparison between a region map and a randomly generated voronoi map (*p* < 0.001). In summary, regions and domains are similar between recordings either in the same or on different animals, compared to a null map distribution.

### Extracting Optimized Time Courses

At full resolution, there are approximately 1.5 million pixels along the surface of the cortex– an impractical number of sources for most network analyses, which work best on 10-300 time series.^29^ We propose that these data-driven domain maps are an optimal method for extracting time courses from the cortical surface. Time series were extracted by averaging the filtered movie under each domain. This results in a series of 230 ± 14 time series per video recording, representing a ∼ 6,500-fold reduction in size (fig. 4a).

**Figure 4:**
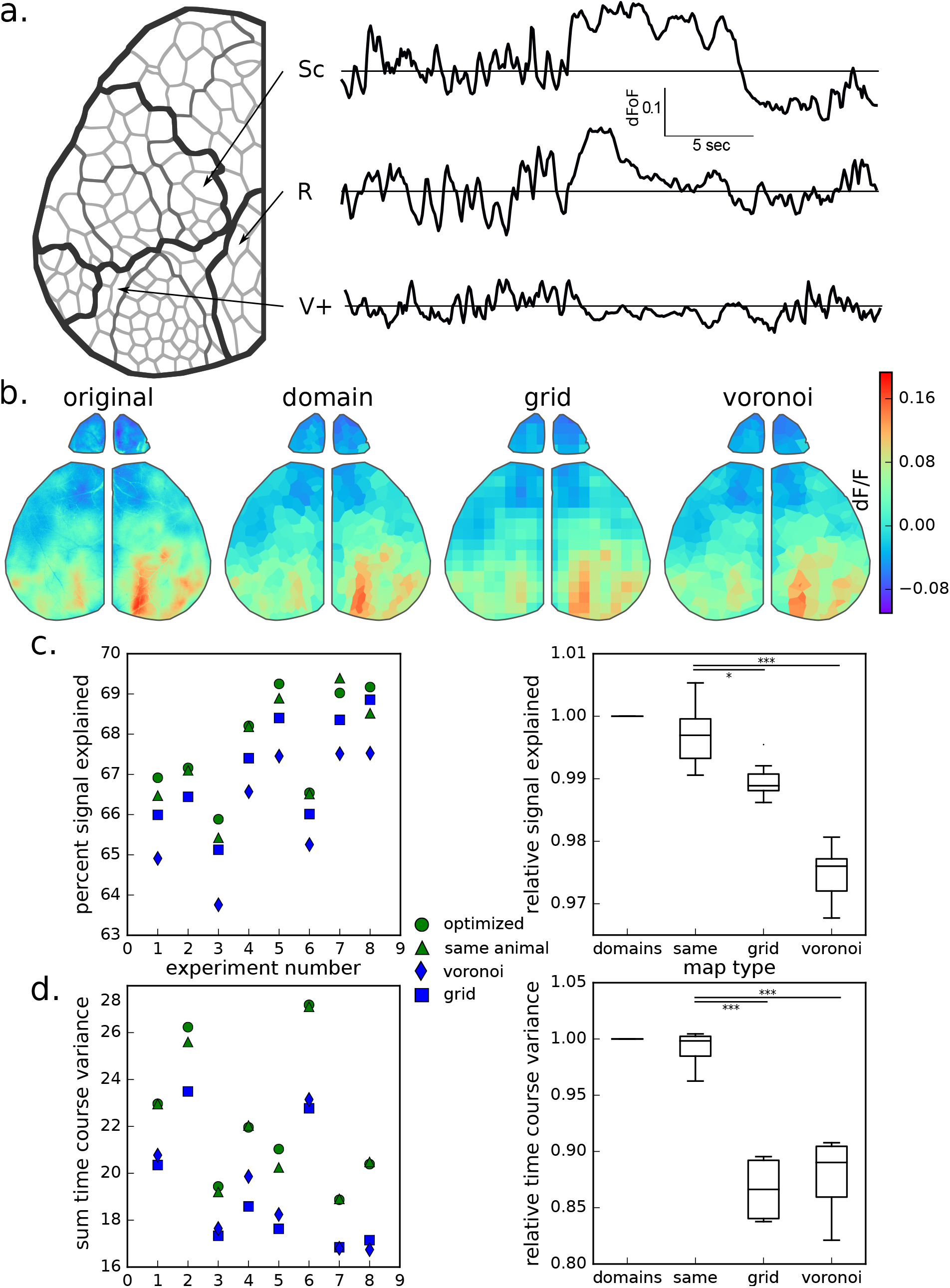
Time series extracted from Domain maps outperform time series generated from other methods. a) Schematic of mean time courses extracted from different domains generated from a domain map. b) Example of a mosaic movie frame rebuilt with respect to each downsampling technique. The filtered movie is replicated in the upper left corner. The upper right image corresponds to the representation of the same video frame rebuilt from the extracted domain time series. The left and right bottom panels correspond to the representation of the same video frame rebuilt from the extracted domain time series from either grid or voronoi maps, respectively. c) Percent total signal of the filtered video represented by extracted time courses. Percent of overall video signal captured in domain maps was calculated from one video from each animal (green circle; n=8), and compared to a map generated from a separate video calculated from the same animal (green triangle). Percent total signal represented by time courses extracted from grid (blue square) or randomly generated (blue diamond) maps were compared as controls. In the right panel, the percent signal relative to the domain map percent signal was summarized in a box plot. d) Variation between time courses extracted with each map method was then quantified as a sum signal variation for each experiment. In the right panel, the sum signal variation for each comparison map relative to the optimized domain map sum signal variation was summarized in a box plot.

To test how well the full filtered video was represented in these time series, we rebuilt ‘mosaic movies’, where each domain is represented by its mean extracted signal at any given time point (fig. 4b, fig. S5 (Video)). Comparing the borders of the large higher order visual activation, one can see visually that the data appears more distorted in the voronoi and grid. To numerically compare whether this method of time course extraction was superior to alternate methods, we also compared mosaic movies rebuilt with either grid or voronoi maps.

The residuals between the mosaic movies and the filtered movies were compared to the total spatial variation in the filtered movie to quantify the amount of total signal represented by the extracted time courses (fig. 4c, left). In nearly every experiment, the optimized domain map performed better than any other time course extraction method, and accounted for 68 ± 1.2 % of the total spatial signal in the filtered video (n=8).

Domain maps generated from different videos from the same animal performed nearly as well 6 as the optimized domain maps created from the video compared (fig. 4c, right). These maps performed significantly better (*p* = 0.01) than the grid maps, and much better than the voronoi maps (*p* < 0.001).

Compared to saving the full ICA compressed dataset, saving these extracted time courses and all associated metadata results in a file size of ∼ 100MB, for a ∼ 60-fold additional compression. One potential benefit to accounting for the underlying regions of the brain while extracting time courses is reducing the amount of times that an extracted mean signal is diluted by signal from a neighboring region. Properly restricting time series extraction to statistically independent units should enhance the dynamic range between extracted time series.

To test whether domain maps extracted time courses better extract the full range of variation in the cortical surface, we compared the total variation between time courses rebuilt under domain maps from the same video, same animal, or control grid and voronoi maps (fig. 4d, left). When normalized to the performance of the optimized domain map, domain maps from the same animal again had similar performance, but grid and voronoi maps performed significantly worse (*p* < 0.001; fig. 4d, right). There is a ∼ 15% reduction in signal variation in grid or voronoi maps compared to domain map extracted time courses.

## Discussion

Here we have shown that high resolution imaging of mesoscale cortical calcium dynamics combined with data-driven decomposition using ICA results in an optimized extraction of neural source signals. We demonstrate that these methods provide precise isolation and filtration of video artifacts due to movement, optical deformations, or blood vessel dynamics while recovering cortical source signals with minimal alteration.^27^ This approach can either be used alone, or in conjunction with techniques to correct calcium dynamics from tissue hemodynamics.^10, 20, 30^

Signal separation from mesoscale calcium dynamics recorded across the cortical surface was the most complete at the highest spatial resolution tested (pixel size of 6.9 µm/px). Temporal resolution had less of an effect on ICA signal separation; we found that a 10Hz sampling rate was sufficient. These metrics for signal quality are automatically generated by our algorithm, and can be used to optimize signal collection on any given experimental setup. The number of components identified is highly stable after recording sufficient duration of dynamics, and provides a metric for spatial complexity of neural signal across the neocortex. Compared with the high density optical recordings we used here, other neurophysiological techniques are limited in the number of available spatial samples. Thus the effect of signal recording resolution on ICA decomposition of neural signal sources had not previously been reported.

We further demonstrate an ICA-based method for using these components to perform a data-driven mapping of the captured cortical dynamics, resulting in a superior isolation of the various signal sources on the cortical surface. Domain maps generated are specific to individual animals. Detected units vary in shape and size across the cortical surface, and have features that resemble known cortical morphology. These maps could help elucidate changes in functional structure across the cortical surface across different ages groups, or genetic conditions known to change cortical spatial structure.

Using optimized signal extraction results in a higher dynamic range of extracted time series, and thus would produce correlation maps and network analyses with enhanced separation between neighboring regions. Stimulation experiments or anatomical post-processing of brain tissue could provide insights on how well the sorted domain regions correspond with known functional and anatomical units of the brain, and may provide a new framework for unbiased exploration of novel functional domains and interactions. If these maps align well to anatomical regions of interest, they can additionally be used as a reference for in-vivo cortical mapping, with potential applications in live feedback or targeted injection experiments.

An additional benefit is the highly compressed data format. The original or video can be rebuilt with relatively little loss of information from a reduced set of ICA components, for a ∼ 10x reduction in file size. This results in a representation of a high resolution video that is much more manageable to work with on a local computer.

The methods presented here address the most common issues in analyzing large wide-field mesoscale datasets, including filtration of vessel artifacts, spatial mapping, and optimized time series analysis. With these tools, neuroscientists can easily collect and analyze high quality neural dynamics across the cortical surface, allowing the investigation of complex networks at unprecedented scale.

**Figure S1:**
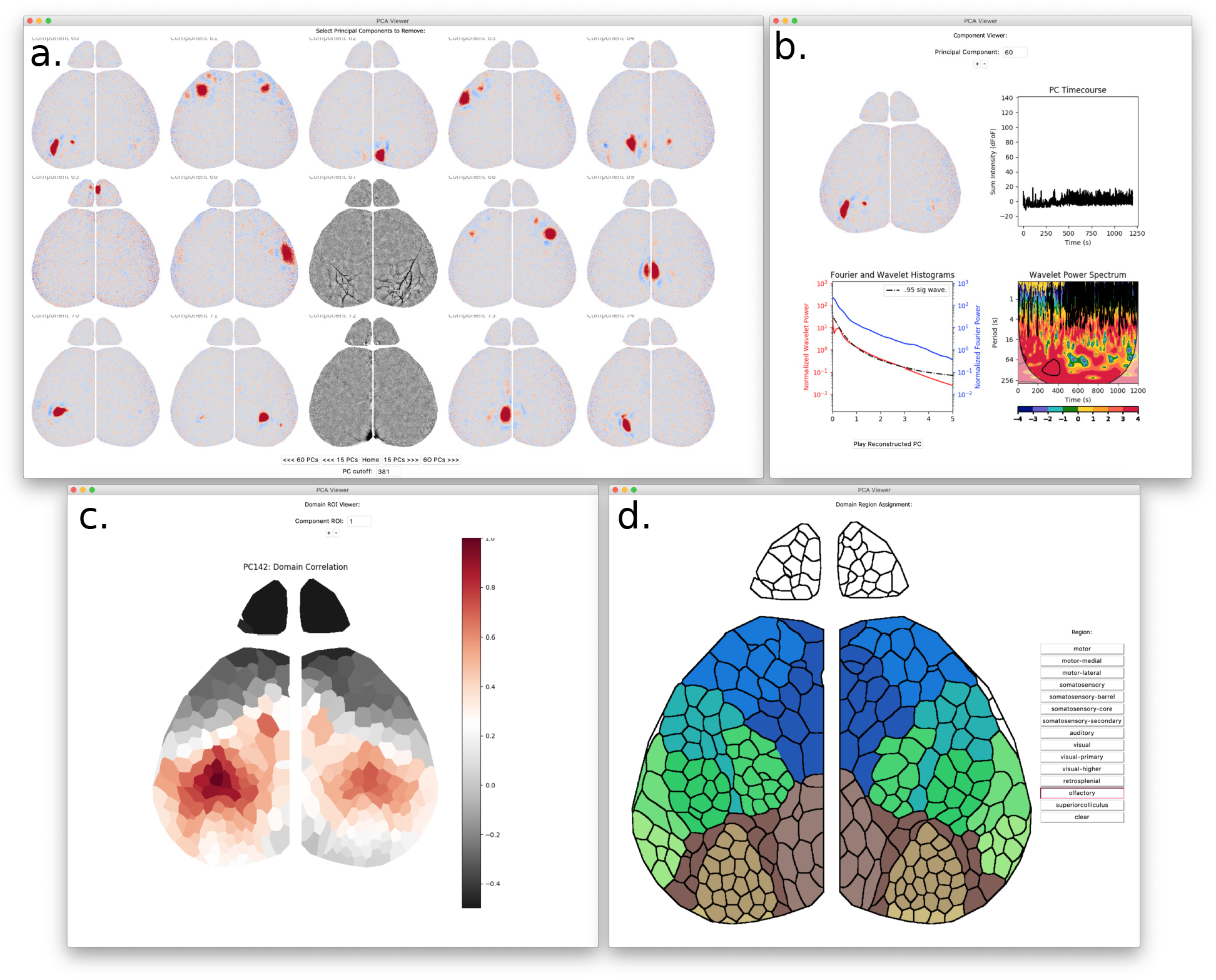
A Tkinter-based graphical user interface (GUI) for browsing independent component analysis results. a) 15 independent components, order 60-74 by variance. Components displayed in grey are selected as artifact either manually or using a machine learning classifier. A click on the display for any given component manually toggles its classification as either signal or artifact associated. Components colored in the cool/warm colormap are signal associated. Components colored in the black/white colormap are artifact associated. Buttons on the bottom panel control GUI movement through the dataset. The text panel at the bottom displays where the index for the signal/noise cutoff. b) The component viewer displays additional temporal metrics about any given component. The top controls allow movement through the dataset by manual scrolling with (+/-) buttons, up/down keys, or through typing a desired component in the text box. PC timecourse displays the mixing matrix timecourse extracted by ICA for the given components. The Wavelet power spectrum is displayed in the bottom right, and an integrated wavelet or fourier representation is available on the bottom left. 0.95 significance as estimated by the AR(1) red-noise null hypothesis is displayed as a dot-dash line. c) The domain map correlation page shows the pearson cross correlation value between a selected seed domain and every other domain detected on the cortical surface. The seed domain can be changed through the arrow keys, the (+/-) buttons, or by clicking on a different domain on the displayed domain map. d) The Component region assignment page allows manual region assignment for each domain. After the region is selected from the menu on the right, each domain clicked on the domain map is assigned to that region.

**Figure S2:**
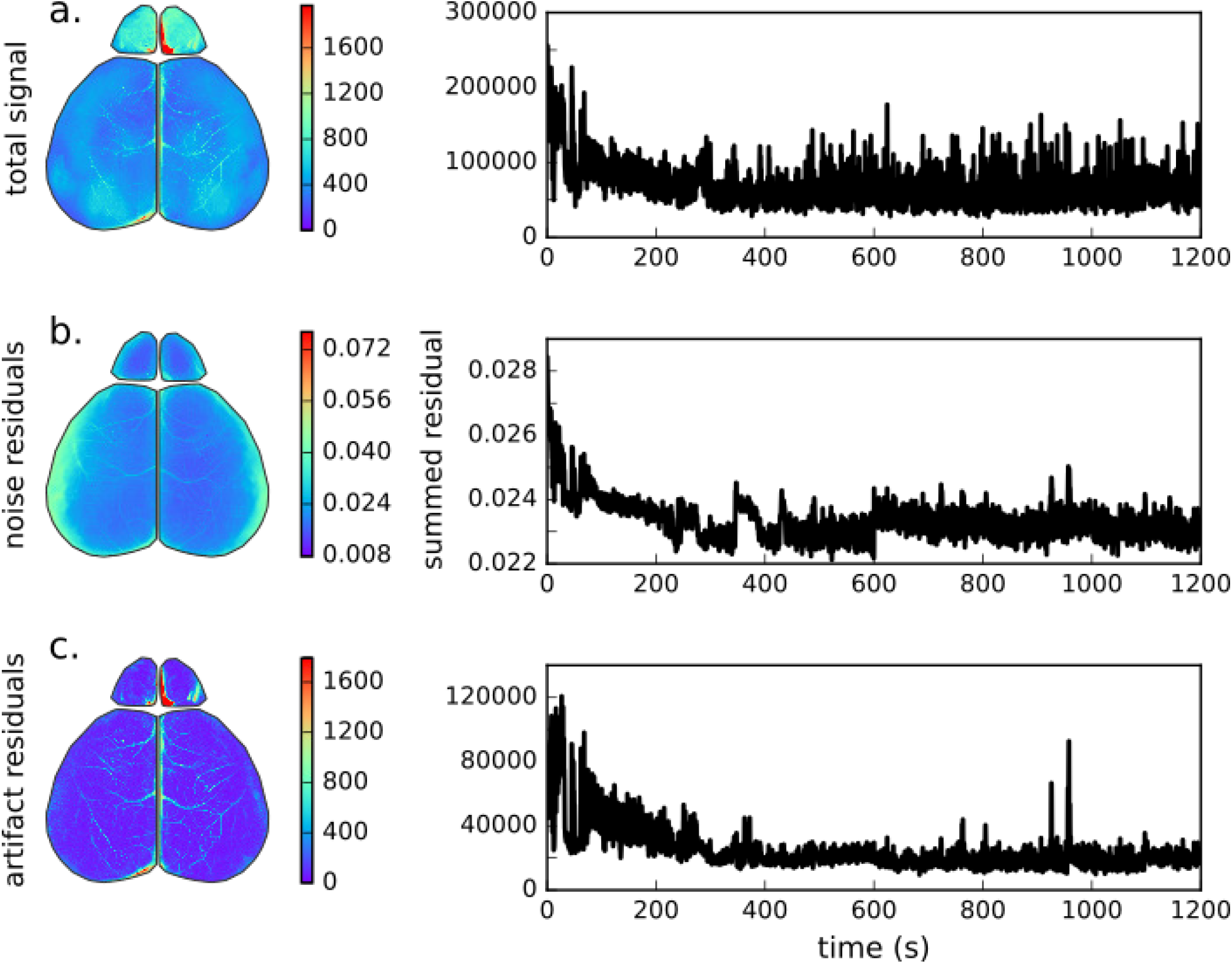
Comparison of spatial and temporal information content through compression and filtering. a) The original spatial information captured as quantified by a mean subtracted absolute value projected spatially (left) or temporally (right). b) The difference in information between the original input data and the rebuilt ICA projection, excluding noise components beyond the 25% saved in the processed file. The difference movie is projected spatially or temporally to visualize where information was lost in compression. c) Information removed by artifact filter. The artifact movie is rebuilt and projected spatially or temporally to visualize where information was modified by the ICA-based artifact filter.

**Figure S3:**
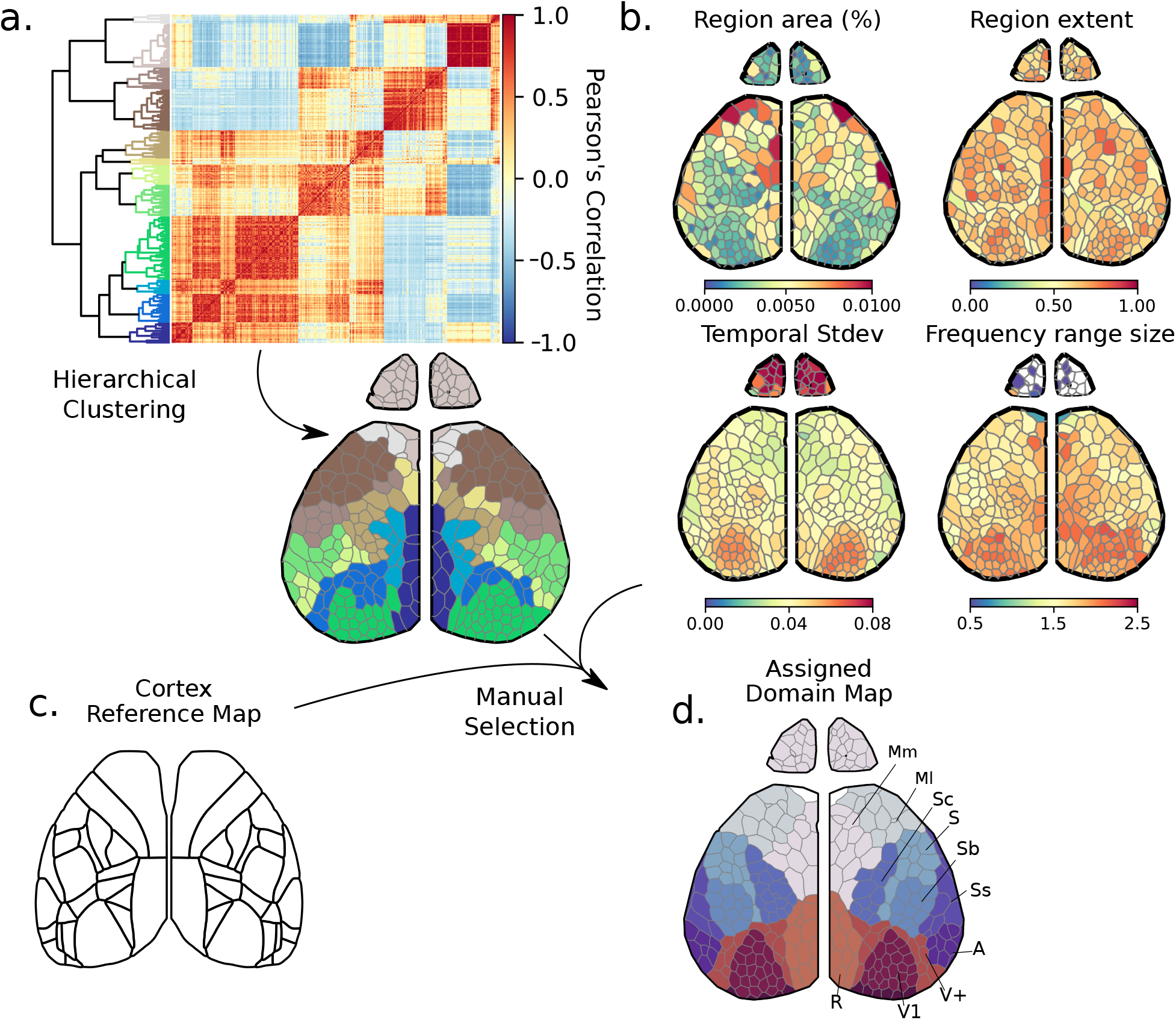
Using data-guided methods to assign domains to cortical regions. a) Hierarchical clustering based on Pearson’s correlation produces a set of ∼ 13 regions across the cortical surface. (b) Domains colored by various calculated spatial and temporal metrics to aid region assignment. Region area is calculated as a percent of the total cortical surface. Region extend ranges from 0 to 1 and calculates the relative area of a domain to its bounding box. Temporal standard deviation is calculated from the extracted time series, and frequency range size is calculated from wavelet significance. c) The Allen Brain atlas map^31^ is additionally used for anatomical reference. d) The final manually assigned region, with associated labels.

**Figure S4 (Video):**
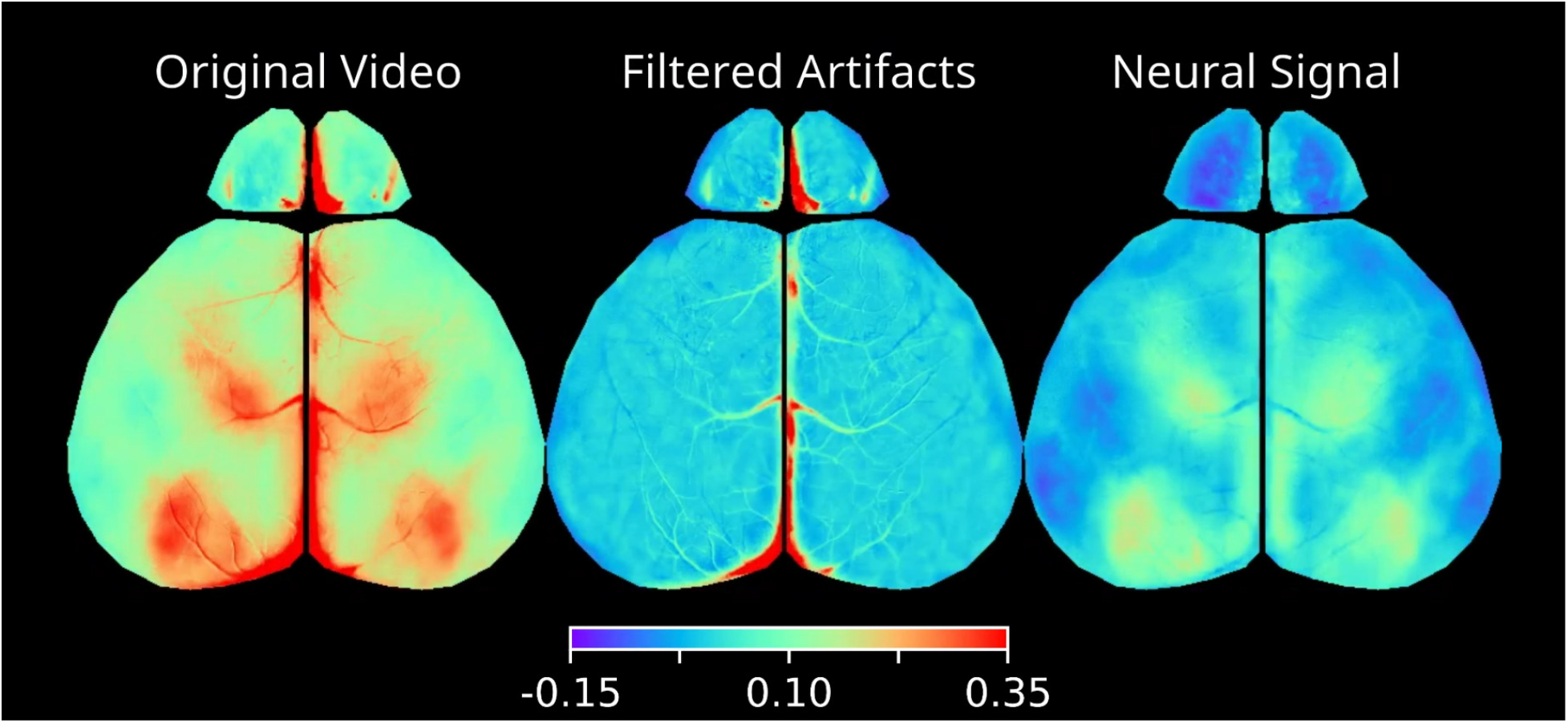
ICA filtration removes artifacts for superior neural signal unmixing. Original video (left) is decomposed into artifact components and neural signal. The filtered artifact movie (center) can be rebuilt to visualize artifacts that were isolated and removed during the filtration process. The rebuilt neural signal (right) depicts just the filtered neural signal. 0.5Hz filtered mean is re-added to both filtered artifacts and neural signal (^27^). Video is a real-time 1-minute excerpt. Values displayed are in dF/F.

**Figure S5 (Video):**
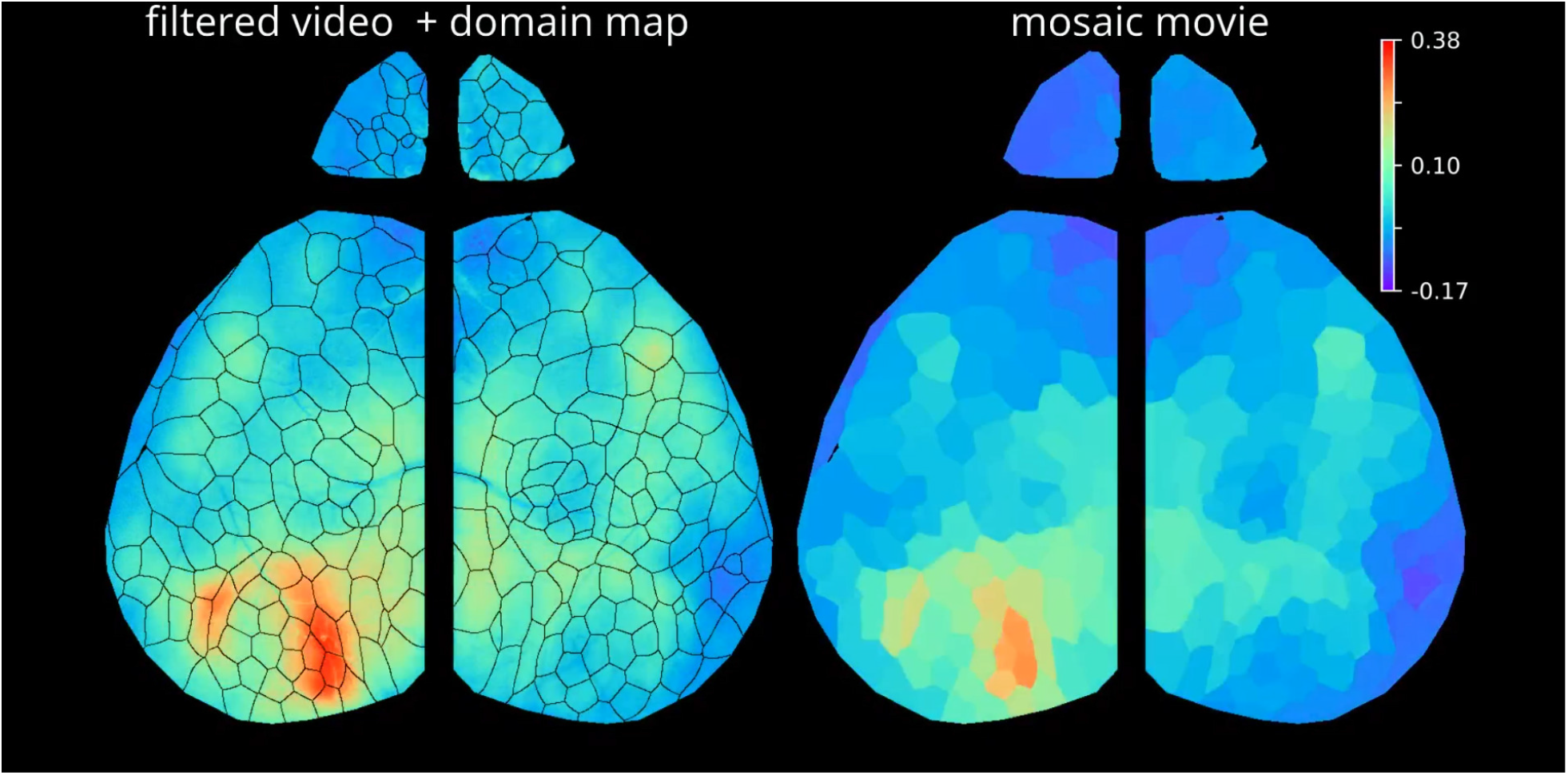
The mosaic movie represents the neural signal captured from time series extracted from domains across the cortical surface. Signal from the filtered video (left) is segmented by the data-driven domain map (left overlay). Average time series from these segmented domains can visualized as a mosaic movie, where each domain is represented by its averaged time series. The filtered video contains 1.77 megapixels representing the cortical signal, while the mosaic movie contains only 300 unique time series to describe the same signal with a 5900x compression rate. Video is a real-time 1-minute excerpt. Values displayed are in dF/F.

## Supporting information

Figure S4: ICA filtration removes artifacts for superior neural signal unmixing.

Figure S5: The mosaic movie represents the neural signal captured from time series extracted from domains across the cortical surface.

## Abbreviations and Terms Defined

ΔF/F (dFoF): change in fluorescence over mean fluorescence
ICA: Independent Component Analysis
PCA: Principal Component Analysis
Domain Map: maximum projection map of ICA components
Domain: A single contiguous unit from a domain map, represents an ICA component’s maximal region of influence
Mosiac Movie: a video representation of the time series extracted under each domain in the domain map

---

**Region Abbreviations** (Used for Region Maps in fig. 3).

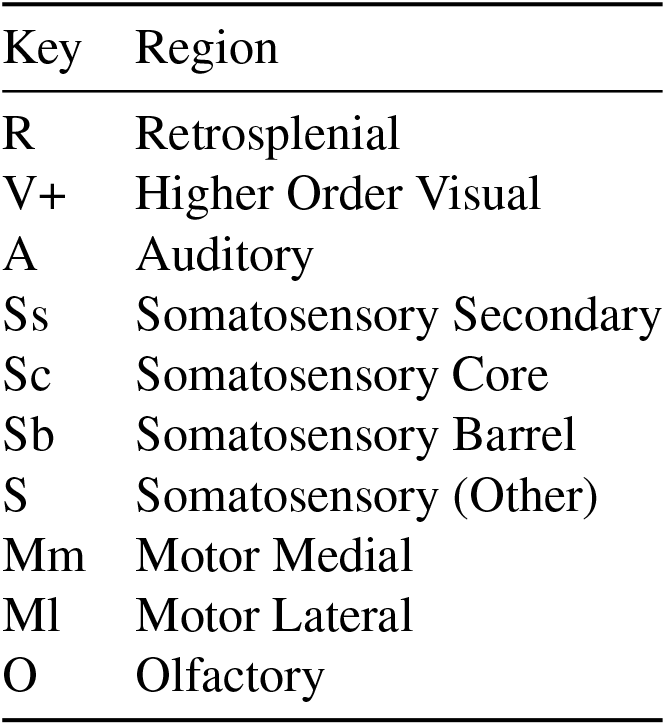

## Methods

### Mice

All animal studies were conducted in accordance with the UCSC Office of Animal Research Oversight and Institutional Animal Care and Use Committee protocols. P21-22 Snap25 GCaMP6s transgenic mice (JAX: 025111), were maintained on a C57/Bl6 background in UCSCs mouse facilities. To identify Snap25 GCaMP expressing mice, a single common forward primer (5’-CCC AGT TGA GAT TGG AAA GTG-3’) was used in conjunction with either transgene specific reverse primer (5’-ACT TCG CAC AGG ATC CAA GA-3’; 230 band size) or control reverse primer (5’-CTG GTT TTG TTG GAA TCA GC-3’; 498 band size). The expression of this transgene resulted in pan-neuronal expression of GCaMP6s throughout the nervous system. At the end of each recording session, the animal was euthanized and the brain dissected.

Quantifications in ICA-based Filtering and Segmentation was performed on 4 sets of Snap25-GCaMP6s littermates. These animals included experimental and control mice a study looking at microbiota influence on cerebral networks.^32^ These methods work independent of experimental condition and the microbiota had little effect on domain parcellation.

### Surgical procedure

All mice were anesthetized with isoflurane (2.5% in pure oxygen) for the procedure. Body temperature was maintained at 37C for the duration of the surgery and recovery using a feedback-regulated heading pad. Lidocaine (1%) was applied subcutaneous on the scalp, followed by careful removal of skin above the skull. Ophthalmic ointment was used protect the eyes during the surgery. The head was glued using cyanoacrylate to two head bars, one across the back of the skull and the other on the lateral parietal bone.

After the surgery was complete, mice were transferred to a rotating disk for the duration of the recording. At the end of the recording session, the animal was either euthanized or perfused and the brain dissected.

### Recording calcium dynamics

In-vivo wide-field fluorescence recordings were collected in a minimally invasive manner. Imaging through the skull by single-photon excitation light from two blue LED light (470 nm, max power 1000 mW; Thorlabs M470L3) produces a green fluorescent signal that is collected through coupled 50*mm* Nikon lenses (f5.6 / f1.2, optical magnification ∼ 1x) into a scientific CMOS camera (PCO Edge 5.5MP, 6.5*µ*m pixel resolution). The top lens in the tandem lens setup was used to focus on the cortical surface, thereby lowering the magnification slightly; anatomical representation for each pixel corresponded to 6.9 ± 0.2*µ*m (min: 6.7*µ*m, max: 7.2*µ*m). Excitation light was filtered with a 480/30 nm bandpass (Chroma Technology AT480/30x) and the emission signal was filtered with 520/36 nm bandpass (Edmund Optics 67-044). Data collection was performed in a dark, quiet room with minimal changes in ambient light or sound. Raw data was written directly as a set of 16 bit multi-image TIFF files.

All video segments consisted of a set of continuously collected images at 10 or 20 frames per second for 10 minutes. The total amount of recorded data for each animal was generally at least 40 min and the amount of time in between video segments was less than 1 minute. When more than 10 minutes of video data was used for single decompositions, multiple videos were concatenated together. All analyses were conducted on data recorded at 10Hz, except exploration of effects of resolution on data quality.

Spatial resolution analyses were performed on a single 10 minute recording at 10Hz. Spatial down sampling was conducted by taking the mean between groups of pixels in a dxd square, where d is the integer downsampling factor. Temporal resolution analyses were performed on a single 10 minute recording at 20Hz. This data was temporally downsampled by taking the mean between subsequent frames.

### ICA decomposition and saving

ICA was performed using FastICA,^33^ implemented through python’s sklearn decomposition.^34^ The ICA decomposition was applied to the spatially flattened (xy,t) 2-D representation of the video data under the cortical ROI mask. The mean time series is pre-subtracted from the array before SVD decomposition or ICA decomposition, since ICA cannot separate sources with a mean signal effect. The filtered, unfiltered mean, ICA components, mixing matrix, and associated metadata are all saved. Data is stored and saved in this flattened format for storage optimization. Components are locally spatially reconstructed for visualization in the GUI.

Requesting the full number of components resulted in extremely lengthy processing times. To reduce the processing time, the data was preprocessed through Singular Value Decomposition (SVD) whitening, and noise components were cropped. To ensure that no signal was lost, and there were ample dimensions left for ICA separation, the inflection point between SVD signal and noise floor was identified, and SVD components were reduced to include components equal to 5 times the SVD signal to noise cutoff value. This multiple cutoff can be adjusted while ICA projecting.

After calculating and sorting the ICA results, excessive noise components are removed from the dataset for compression. The cutoff was determined by identifying the inflection point in the lag-1 autocorrelation distribution with a two-peaked KDE fit. Components were saved such that 75% of the components saved were signal or artifact, and 25% of the components saved were noise associated. If not enough noise components were returned by the ICA decomposition, there is a risk that signals were not sufficiently unmixed, so the ICA decomposition was repeated with a higher SVD cutoff until enough additional noise components were included.

For resolution downsampling analysis, the number of components requested was kept constant for all resolution values to avoid confounding the results. The cutoff for the highest resolution tested was used for all lower resolution decompositions.

ICA returns components that are unsorted and often flipped. Components were first sorted by their time series variance. The spatial histogram of eigenvector values across each component can be visualized as a single tailed gaussian distribution centered around 0, where the tail represents the spatial domain of each component. The two edges of the distribution are first identified. The boundary closer to 0 is taken as the edge of the central noise distribution, and that boundary is used to define the dynamic threshold on the opposite side of 0. Any values outside of this noise distribution and is part of the wider tail are included in the binarized domain of the component. If the tail was negative, the component was flipped spatially and temporally. In this way, components were all identified as positive affectors for visualization purposes. Movie rebuilding was not affected by this process.

All code used in this paper will be available at github.com/ackmanlab/pyseas or as a package (pySEAS: python Signal Extraction and Segmentation) in the python package index (`pip install seas`).

### Data processing

ICA decompositions of videos at full spatial resolution and duration (20 min) were run on a single cpu node of a computing cluster having 1024 GB of RAM. Shorter recording length videos (10 min) could be processed on a node having 512 GB of RAM. After ICA processing, map creation and time series analysis were performed on local computers having 16-32GB of RAM.

### Map creation and comparisons

Domain maps were created by separating the cortex into regions represented by different ICA components. Each component was blurred by a 51-pixel kernel, then the maximum projection was taken through the component layers. The resulting data is a cortical map that denotes the component with maximum influence over any given pixel.

This map was then further processed to get rid of domains smaller than 1/10th the mean domain. Any domain smaller than this size is checked to see if the second most significant component would produce a larger continuous structure. If after a few loops of this, pixels cannot be assigned into a larger structure, the points are excluded from the final map. Indices are then adjusted such that any non-continuous regions represented by the same domain are assigned to different units.

Olfactory bulbs were included in map generation, but domains were highly variable, and were excluded from map quantifications.

Voronoi maps were created to match the same number of domains or regions as the original map, *n*, and shares the same cortical mask as the original map. To create this map, *n* points were distributed randomly across the cortical mask. To turn these points into regions, the voronoi diagram was created using the scipy spatial package^35^ and was applied as a voronoi map.

Grid maps were created to match or exceed the same number of units as the original map, and share the same cortical mask as the original map. A uniformly spaced 2D grid was placed over the original map, and resulting units were counted. If the number of resulting spatial units exceeded that of the original map by < 15, the map was accepted as a valid comparison. Otherwise, the map was rejected and a new grid map was calculated.

For every domain or region in the original map, the nearest neighbor was identified in the comparison map with a KNN tree. To quantify the spatial similarity of each identified domain or region, the Jaccard index (spatial overlap / union) was then calculated. For each comparison, *n* Jaccard indices were calculated.

When comparing maps generated from different animals, the optimal alignment was calculated by shifting the second map up to 100 pixels in any direction. The optimal direction was determined by maximizing the Jaccard overlap. Each generated map was compared to one map from the same animal, one littermate, and two non-littermates, as well as one randomly generated voronoi map.

### Compression and filtering residuals

Compression residuals are calculated while saving the ICA decomposition results. The original movie is rebuilt from the reduced ICA results, and residuals are calculated by taking the absolute value of the difference between the two videos. The spatial and temporal projection of this absolute difference movie is saved as the spatial and temporal residuals of the decomposition, and is stored as metadata with each ICA decomposition.

### Domain residuals and domain signal analyses

To quantify the amount of signal present in the original movie that was not included in the domain map, residuals were calculated by subtracting the ‘mosaic movie’, representing time series from each spatial domain from the original movie. The absolute value was then applied so that all numbers represented a positive difference, and residuals were summed to create a single value. The time series was not re-added to either the original movie or mosaic movie, since this can be easily summarized as a different temporal metric. To represent the amount of relative variation to the original dataset, this number was compared to the summed absolute value of the mean-subtracted original movie.

### Statistical significance

Statistical significance was calculated using OLS models from `statsmodel.formula.api` with Holm-Sidak multiple testing correction (*p* ≤ 0.5: *; *p* ≤ 0.01: **; *p* ≤ 0.001: ***). Model significance is determined by the F-statistic, and significance of two-group analyses (p>|t|) are calculated with t-tests.

## Acknowledgements

The authors acknowledge C. Santo Thomas for maintaining the lab mouse lines, and University of California Santa Cruz’s Hummingbird Computational Cluster for support and node maintenance. This work was supported by Startup funds from University of California, Santa Cruz, Division of Physical and Biological Sciences, grants from the National Institutes of Health, USA (NIH T32 GM 133391) to S.C.W. and B.R.M, and by a Hellman Fellows Fund Award to J.B.A. Funding for D.A. was provided by Maximizing Access to Research Careers (MARC) and Initiative for Maximizing Student Development (IMSD).

## Contributions

ICA filtering, exploratory GUI, map creation and time series extraction and analysis code, was written by S.C.W. All recordings, metric extractions, mean frequency analysis, and domain assignments were performed by B.R.M. Control wavelet analysis was assisted by D.A. J.B.A. oversaw the project and provided feedback to experimental design, results, and paper preparation. The manuscript was prepared by S.C.W, with input from all authors.

